# Targeting “Immunogenic Hotspots” in Dengue and Zika Virus: A Novel Approach to a Common Vaccine

**DOI:** 10.1101/2021.07.23.453561

**Authors:** Dhrubajyoti Mahata, Debangshu Mukherjee, Vanshika Malviya, Gayatri Mukherjee

**Author notes:** To whom correspondence should be addressed: Gayatri Mukherjee, School of Medical Science and Technology, Indian Institute of Technology Kharagpur, West Bengal - 721302, India. These authors have contributed equally to the work and share first authorship (Dhrubajyoti Mahata, Debangshu Mukherjee).

## Abstract

Diseases caused by Dengue (DENV) and Zika (ZIKV) viruses cause significant mortality and illness globally. Due to the high sequence similarity of the viral proteins and the purported cross-reactive immune responses against the viruses, we envisioned a common multi-epitope vaccine (MEV) against both viruses by adopting a novel approach of identifying “immunogenic hotspots”. These stretches of the structural and non-structural proteins are enriched with MHC class I and class II supertype-restricted T cell epitopes, and B cell epitopes, in addition to being highly conserved between different DENV serotypes and ZIKV. Such an approach ensures inclusion of multiple overlapping T and B cell epitopes common to both viruses, and also warrants high population coverage. Importantly, epitopes known to cause antibody-dependent-enhancement of infection have been excluded. These immunogenic hotspots have then been stitched together with linkers in-silico along with an adjuvant, CTxB to develop the MEV candidate. Four structural models of the MEV were selected on the basis of conformational preservation of CTxB, and their biophysical parameters, which also conserved the immunogenicity of the multiple epitopes. Importantly, each of the MEV candidates were found to interact with TLR4-MD2 complex by molecular docking studies, indicative of their ability to induce TLR-mediated immune responses.

## Introduction

Vector-borne viral diseases, like Dengue fever and Zika viral fever cause severe illness, significant mortality and resultant socio-economic burden worldwide, thus proving to be a major public health concern of the modern era. The Dengue virus (DENV) has four serotypes (DENV1-4), which are responsible for dengue fever, dengue hemorrhagic fever, and dengue shock syndrome in humans[1]. Globally, it is the fastest-growing mosquito-borne disease, causing nearly 400 million infections every year with a mortality rate of 20% when untreated (https://www.who.int/en/news-room/fact-sheets/detail/dengue-and-severe-dengue). On the other hand, the Zika virus (ZIKV) causes the zika viral fever which, in addition to causing headache, muscle and joint pain, rash, edema of extremities, retro-orbital pain and conjunctival hyperemia [2],often lead to congenital abnormalities in unborn fetus [3]. Due to such long-lasting damage by ZIKV, WHO declared this virus a public health emergency of international concern in 2016.

Being members of the genus Flavivirus, DENV and ZIKV share some common genetic as well as phenotypic features. Both are borne by different species of the *Aedes* mosquito, and have approximately 11 kb+ ssRNA genome that encodes three structural and seven non-structural proteins. The three structural proteins are capsid (C), envelope (E) and the membrane glycoprotein (M); while the seven non-structural proteins are called NS1, NS2A, NS2B, NS3, NS4A, NS4B, NS5[4,5]. Importantly, the 4 serotypes of DENV share a high genetic similarity with ZIKV, especially in the E protein and the NS3 and NS5 proteins[6].

The B cell [7,8] and T cell [9,10,11]mediated immune response against different DENV serotypes show significant cross reactivity not only among themselves but also with ZIKV and vice versa. Such cross-reactive immune response often leads to Antibody Dependent Enhancement (ADE) of infection and disease virulence in successive natural infections with different serotypes of DENV following the first infection. Additionally, ADE has also been reported in ZIKV infections following prior DENV infections and vice versa [12,13,14]. This could be due to binding of the non-neutralizing or weakly neutralizing, but cross-reactive antibodies from the first infection, to the heterotypic virus particles, which could lead to increased viral uptake by monocytes and macrophages via their Fcγ receptors, thus resulting in increased viral load [15,16].Reports have shown that Dengvaxia, the only licensed vaccine against DENV currently available, poses considerable risk of ADE of infection upon encounter with dengue serotypes in vaccinated individuals who were seronegative at the time of vaccination [17,18,19]. Therefore, it is imperative to design and develop a new generation vaccine which will be able to protect against all DENV serotypes and ZIKV, while circumventing the deleterious effects of ADE.

In the current study, we first identified the viral proteins from all four DENV serotypes and the Asian strain of ZIKV that have significant antigenic probability and phylogenetic relatedness. These proteins were then probed for T cell epitopes restricted to MHC class I and class II supertypes, instead of individual MHC alleles, thereby ensuring a high population coverage, to account for the high degree of polymorphisms in the HLA alleles. Next, we proceeded to map the T cell epitopes on the selected proteins, and identified amino acid stretches of these proteins that- a) contained multiple T cell epitopes restricted to multiple HLA supertypes; b) was present on all four DENV serotypes as well as ZIKV; and c) was highly conserved among the selected DENV strains and ZIKV. These amino acid stretches were henceforth referred to as “immunogenic hotspots”. The hotspots were further probed for the presence of linear and conformational B cell epitopes that were completely conserved in all five viral strains. However, we excluded the immunogenic hotspots that contained B cell epitopes previously reported to be responsible for ADE among DENV serotypes and/or ZIKV, to minimize the deleterious side effects of the vaccine candidate. With this strategy, we identified four completely conserved, multi-epitope rich immunogenic hotspots, which were then joined together with linkers in silico along with CTxB, a well-established adjuvant used for multi-epitope vaccine (MEV) design. Three-dimensional structures of the MEV were predicted with different order of the immunogenic hotspots, using trRosetta server(https://yanglab.nankai.edu.cn/trRosetta/). Models with the highest structural preservation of the CTxB adjuvant were further selected, followed by calculation of the free energy of folding of each model. Thereby four MEV models have been predicted, which were also found to interact with TLR4-MD2 complex by molecular docking studies. To the best of our knowledge, this work represents the first attempt to utilize conserved immunogenic hotspots in the viral protein sequences, to engineer a common MEV candidate with the potential to protect against DENV serotypes and ZIKV, with high population coverage.

## Methods

### Sequence retrieval

The protein sequences for all the four serotypes of DENV (DENV1, DENV2, DENV3 and DENV4) together with the Asian lineage of ZIKV were retrieved in FASTA format from ViPR (Virus Pathogen Resource Database, https://www.viprbrc.org/brc/home.spg?decorator=vipr). These include the structural proteins, viz. Membrane Glycoprotein (M), Membrane Glycoprotein Precursor (PreM), Anchored Capsid Protein (AncC), Capsid Protein (C) and Envelope Protein (E); and seven non-structural proteins for all the above-mentioned strains. The GenBank accession numbers are KJ755855 (DENV1), KY427085 (DENV2), KU216209 (DENV3), KX845005 (DENV4) and LC21972 (ZIKV).

### Determination of antigenicity

Next, the antigenicity of all the proteins was evaluated using the VaxiJen 2.0 server (http://www.ddg-pharmfac.net/vaxijen/VaxiJen/VaxiJen.html). Unlike most other prediction methods which are based on alignment of protein sequences to known antigenic proteins, VaxiJen is based on an alignment independent prediction method which takes into consideration the individual properties of the amino acids in the protein sequence[20]. The prediction method takes into account protein classifications, structural and functional similarities between peptides, principal properties of the amino acid itself and those in its neighborhood, to give a more specific and accurate antigenic probability. The protein sequences were fed into the server in FASTA format and the threshold value of antigenicity was set at the recommended cutoff of 0.4, having highest accuracy for viral proteins. The proteins with the antigenicity score more than the set cutoff were considered as most probable antigens.

### Multiple sequence alignment and phylogenetic analysis

The protein sequence of all the 13 proteins common for all 4 DENV serotypes and ZIKV was subjected to multiple sequence alignment (MSA) to find out the conserved sequences. Clustal Omega server (https://www.ebi.ac.uk/Tools/msa/clustalo/)was used for the alignments; this server utilizes pairwise alignment methods followed by sequence clustering to get distance scores. Following this, a guide tree is constructed and further alignments are done by progressive alignment packages to give the final output and a specific phylogenetic tree of the proteins that were analyzed[21]. Boxshade server (https://embnet.vital-it.ch/software/BOX_form.html) was used for the representation of the MSA data. The amino acid at a particular position was considered conserved if present in the protein sequences of four or more viral strains under consideration. The phylogenetic trees were used to analyze the phylogenetic hierarchy between the proteins from the viral strains.

### Prediction of MHC class I and II restricted T cell epitopes

Following the VaxiJen analysis, MSA and phylogenetic analysis; three proteins, (E, PreM and NS5) were considered the most antigenic. The MHC class I and II epitopes were predicted for these proteins using the IEDB server(<u>IEDB.org: Free epitope database and prediction resource</u>). The extremely polymorphic MHC molecules called HLA (human leukocyte antigens) in humans, can be clustered on the basis of their main anchor specificity into 9 groups (in case of MHC class I) and 7 groups (with respect to MHC class II) called supertypes[22,23]. Between the supertypes, global population coverage is found to be nearly 100%. Therefore, mapping supertype based epitopes gives us the maximum opportunity to mine out conserved immunodominant regions that can be useful in designing a common vaccine. The latest IEDB recommended 2.22 prediction method was used for epitope prediction for the nine MHC class I allele supertypes, viz. HLA-A01, A02, A03, A24, B07, B27, B44, B58, and B60. For MHC class II, IEDB recommended prediction method was used to predict epitopes for a predefined set of MHC class II alleles that gives maximum population coverage[24]. The epitope length was set as 9mer and 15mer for MHC class I and class II molecules respectively. The output result was in the form of percentile rank and the lower the rank of an epitope the stronger the binding of that particular epitope. A cutoff of 2.0 percentile rank was used to get the best predicted epitopes which was then used to compute the immunodominant regions of the proteins.

### Prediction of linear and conformational B cell epitopes

Linear B cell epitopes were predicted using the ABCpred server (https://webs.iiitd.edu.in/raghava/abcpred/). The FASTA sequence of each protein was submitted to get the predicted epitopes. The threshold score was set at 0.51 (between 0.1 to 1) and length of epitope was set as 16mer because the server shows 65.93% accuracy with equal sensitivity and specificity at this default set up[25]. The highly specific predicted epitopes with a score >0.75 were selected for further analysis.

For prediction of conformational B cell epitopes, Discotope 2.0 server (http://tools.iedb.org/discotope/) was used. The three-dimensional structures (resolution <5 Å) of the envelope protein, PreM and NS5 protein of different DENV serotypes and ZIKV were retrieved from Protein Data Bank (https://www.rcsb.org/) and fed to the Discotope server. The threshold value is set at the recommended −3.7 cutoff, which corresponds to 75% specificity[26].

### Identification of epitope rich “Immunogenic Hotspots”

The predicted CD4 and CD8 T cell specific epitopes were further mapped on to the selected proteins. Upon doing so, it was found that certain stretches of the amino acid sequences have given a higher number of overlapping epitopes restricted to multiple MHC alleles. Among these, few amino acid stretches were found in all four serotypes of DENV and ZIKV. These common immunodominant regions were referred to as “immunogenic hotspot”. Further, the epitopes obtained from these immunogenic hotspots were mapped back to the consensus sequences generated from multiple sequence alignment to identify the conserved epitope-rich stretches for each selected protein across all viral strains.

Similarly, the predicted linear B cell epitopes were also analyzed to identify epitope-rich immunodominant regions. These immunodominant regions were then further mapped back to the conserved stretches of T cell epitopes. The conserved amino acid stretches that included common T cell and B cell epitopes in all the DENV serotypes as well as ZIKV were identified. These stretches of amino acids are henceforth referred to as immunogenic hotspots, which have the potential to elicit both T cell and B cell mediated immune response, thereby making them lucrative candidates for MEV design.

### Multi-epitopevaccine design

The conserved immunogenic hotspots were further used for the designing of multi epitope peptide vaccine (MEV). We have selected four immunogenic hotspot sequences derived from E and NS5 protein. These four immunogenic stretches were linked together with GPGPGPG linkerswhich minimizes conformational changes in the tertiary structure and provides flexibility to the peptide chains. Peptide adjuvant CTxB (GenBank Id WP_000593519.1), the B subunit of cholera toxin was included in the N terminus of the MEV using rigid EAAAK linkers. The 103 amino acid long functional domain of the CTxB interacts with the GM1 lipid present on the plasma membrane thereby facilitating the entry of the MEV candidate into antigen presenting cells (APC)[27,28]. Further, this N terminal CTxB adjuvant can also activate the TLR mediated immune response by interacting with TLR4[29], thus making it a suitable peptide adjuvant. To predict the most stable structure of the MEV, the four immunogenic stretches were marked as A, B, C & D and joined one after another in all possible combinations keeping the adjuvant fixed at N terminus which gives a total of 24 models, each 198 amino acids long. Furthermore, the tertiary structures of the MEV candidates were predicted using the trRosetta server (https://yanglab.nankai.edu.cn/trRosetta/). trRosetta builds the protein structure based on direct energy minimizations with a restrained Rosetta. The restraints include inter-residue distance and orientation distributions, predicted by a deep residual neural network[30]. Each combination gave one predicted tertiary structure, which was named as model 1 to model 24. They were further aligned with native structure of CTxB (PDB ID 5LZG) using Pymol software (The PyMOL Molecular Graphics System, Version 1.1 Schrödinger, LLC.), and RMSD values were calculated. Models which have aligned more than 80% of the CTxB’s Cα atoms with its native structure and also have RMSD values <1 Å were further selected and checked for stability by calculating Gibbs free energy for folding. The free energy for folding of each model was calculated using the FoldX tool (http://foldxsuite.crg.eu/command/Stability). These parameters are reflective of the overall stability of the predicted protein. Each of the four finally selected MEV models were further docked with TLR4 (PDB ID: 3FXI) using the ZDOCK server version 3.0.2 (https://zdock.umassmed.edu/) [31]. Such interaction would be indicative of uptake of the MEV by immune cells leading to protective responses. The results of docking were visualized using PyMOL software.

## Results and Discussion

### Selection of proteins for epitope prediction

DENV and ZIKV belong to the genus Flavivirus and have common structural proteins consisting of conserved regions. In case of DENV and ZIKV infection, it is observed that neutralizing antibodies are generated against E, M and its precursor PreM, C and anchored capsid protein (AncC)[32,33,34]. Additionally, T cell mediated immune response in DENV and ZIKV infections has been found to be mostly generated against the non-structural proteins[9,35]. In this study, instead of selecting highly immunogenic individual T and B cell epitopes as vaccine candidates, we aimed to identify T and B cell epitope-rich stretches of both structural and non-structural proteins that are highly conserved among the four DENV serotypes and ZIKV, and formulate a multi-epitope vaccine candidate with these conserved “immunogenic hotspots” of the virus proteome. To select the structural proteins, we used the VaxiJen server which identifies the most probable antigens in an alignment independent manner and considers the properties of the amino acid composition of the protein, protein classification, structural and functional similarities of the peptides. VaxiJen recommends a cut-off of 0.4 in order to select the most probable viral antigens. To satisfy our approach of developing a common vaccine against all four DENV serotypes and ZIKV, we needed to consider the antigenicity of the viral proteins in each individual serotype. Accordingly, we first applied the cutoff to each structural protein (E, M, PreM, C and AncC) of all the DENV serotypes and ZIKV individually, in addition to calculating the average antigenic probability, the cut-off for which was set at 0.6. Only the E (Average antigenicity score =0.6091) and PreM (Average antigenicity score =0.684) satisfied both criteria and therefore were selected for further studies (**Table 1**). This is consistent with the observation that envelope protein induces a strong antibody response against ZIKV[34] and all serotypes of DENV[33]. The cleavage of the PreM into functional M protein is not entirely efficient and as a result, 30-40% of mature virions have PreM protein on their surface[36]. This leads to a considerable antibody response being generated against the PreM protein[37]and thus justifies the selection of the PreM protein at this stage.

**Table 1:**
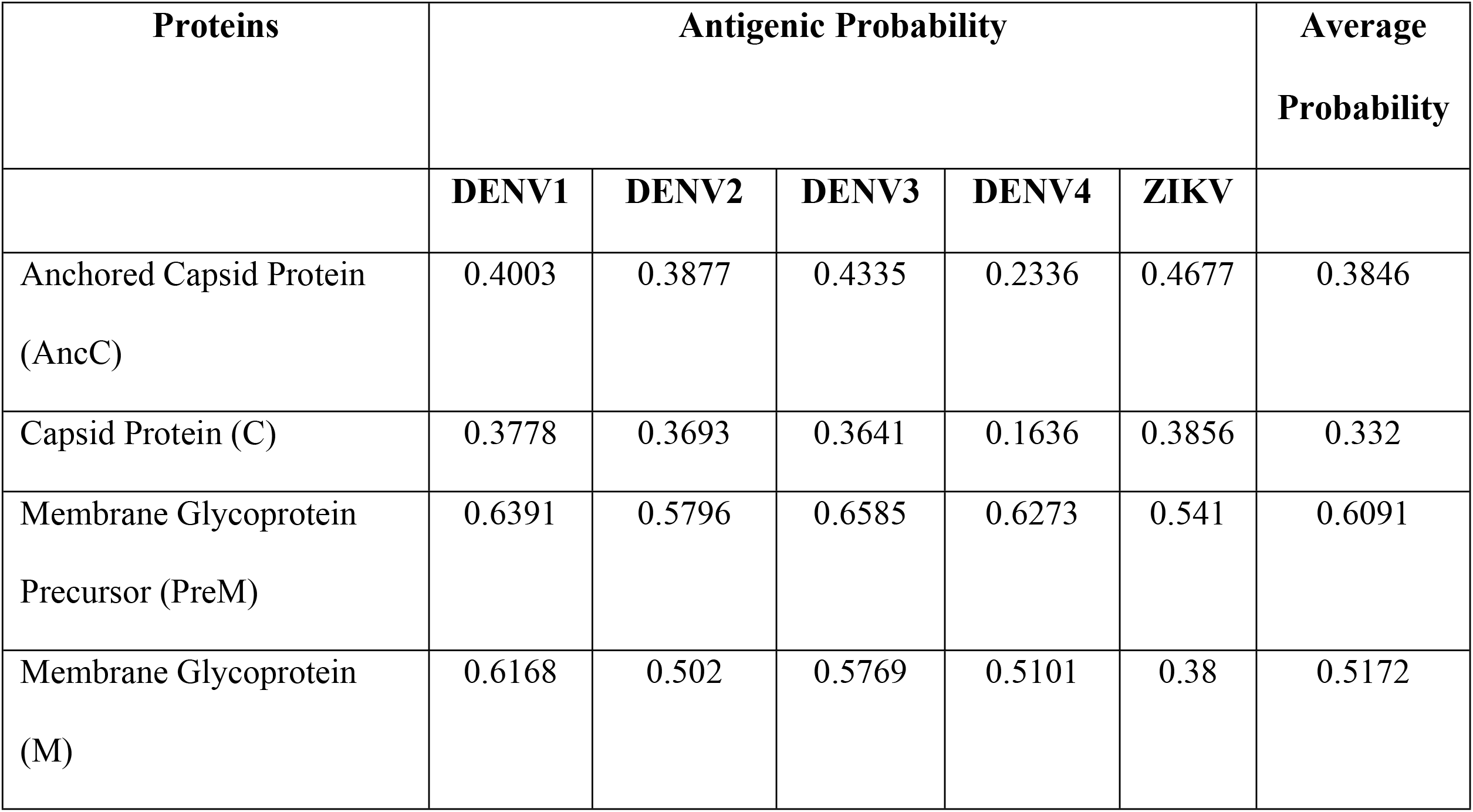
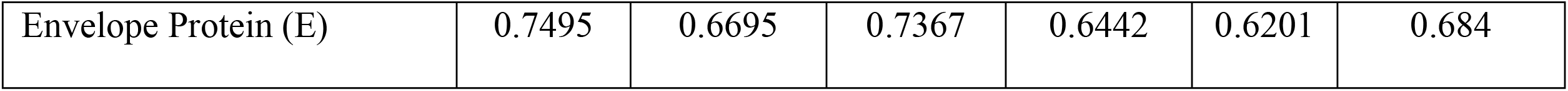
Antigenic probability of the structural proteins as determined by the VaxiJen server.

Like the structural proteins, the non-structural proteins were also investigated on the VaxiJen server to determine their antigenicity. The results showed that all the non-structural proteins qualified the 0.4 cutoff and could be considered probable antigens (**S1Table**). Proteins with high sequence homology and phylogenetic relatedness have greater probability of generating similar or identical epitopes across the species under consideration. In flavivirus, it has been shown that phylogenetic relatedness plays a role in antigenicity, wherein individual distinct clades generate identical epitopes and therefore such relatedness must be considered while predicting the common epitopes[38]. Therefore, to select the best candidates from the non-structural proteins of the DENV serotypes and ZIKV, we probed these proteins for sequence homology determination and phylogenetic relatedness. Our studies as well as already published data [6]show that NS3 and NS5 have the highest sequence homology(greater than 60% homology) between ZIKV and all the DENV serotypes. Therefore, we selected these two proteins for further analysis of phylogenetic relatedness. The NS5 protein of ZIKV and DENV4 share three common ancestors and the pair shares two common ancestors with DENV2, indicative of high degree of relatedness between these three viral species. However, NS5 of DENV1 and DENV3 are phylogenetically separated from these three (**Fig 1(A)**). In comparison, NS3 of ZIKV and DENV4 share two common ancestors, while that of DENV1 and DENV3 share a separate pair of common ancestors, reflective of the greater phylogenetic difference between these two clades. Additionally, further divergence is seen with the NS3 protein of DENV2 from the rest of the members (**Fig 1(B)**). Thus, the data demonstrates that NS5 is more closely related across all the viral species than NS3 and hence it was selected as a candidate for further vaccine design.

**Fig 1:**
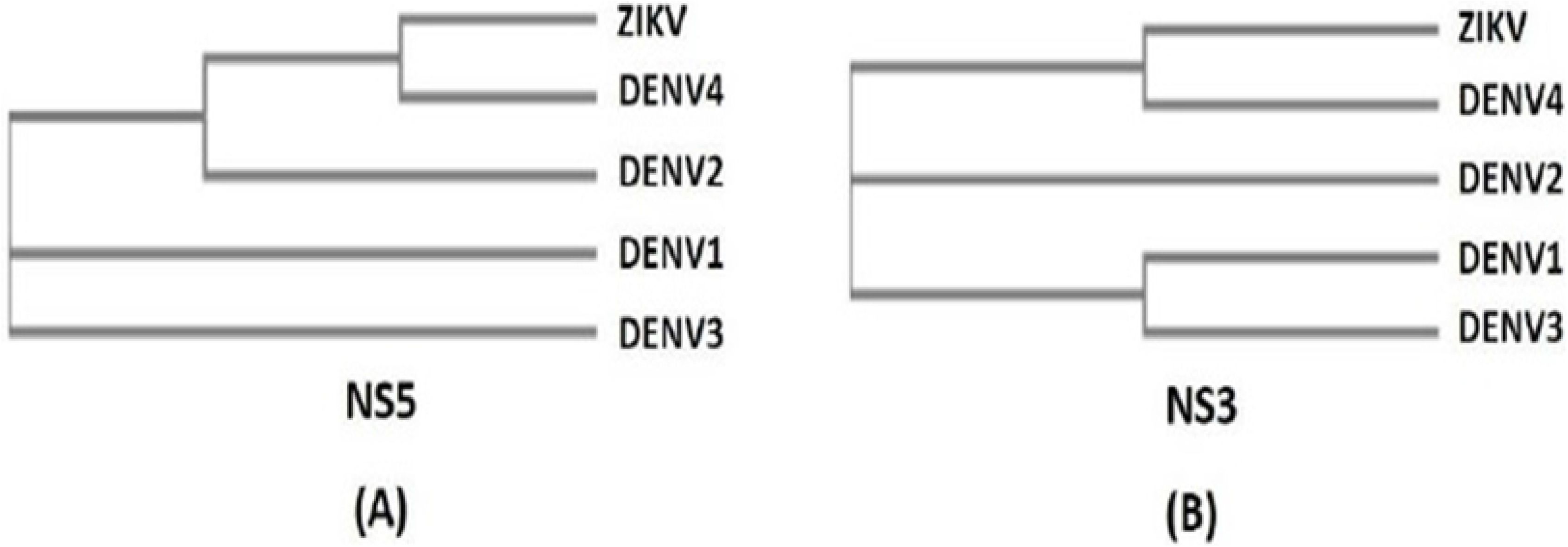
**Phylogenetic tree(A)** NS5 and **(B)** NS3 protein as computed by CLUSTALW server

### Identification of MHC class I restricted epitopes and ‘Immunogenic Hotspots’ from the selected proteins

Based on the antigenicity score and phylogenetic relationship, E, PreM, and NS5 were selected for prediction of MHC class I restricted human CD8 T cell specific epitopes from the IEDB server. The peptide binding cleft of MHC class I can accommodate 8 to 10mer peptides and therefore 9mer peptides were predicted for the nine MHC class I allele supertypes giving maximum population coverage. After analyzing the identified epitopes, it was found that there are distinct amino acid stretches that contain a high number of overlapping epitopes restricted to the different HLA supertypes. These epitope rich regions of the proteins are termed as “immunogenic hotspots”. It was seen that in case of PreM, there was one amino acid stretch that was dominant across all viral species, and contained multiple MHC class I supertype restricted epitopes (**Table 2**). However, upon further analysis, this immunogenic stretch did not yield a conserved peptide stretch that can be used as a hotspot peptide in our MEV candidate.

**Table 2:**
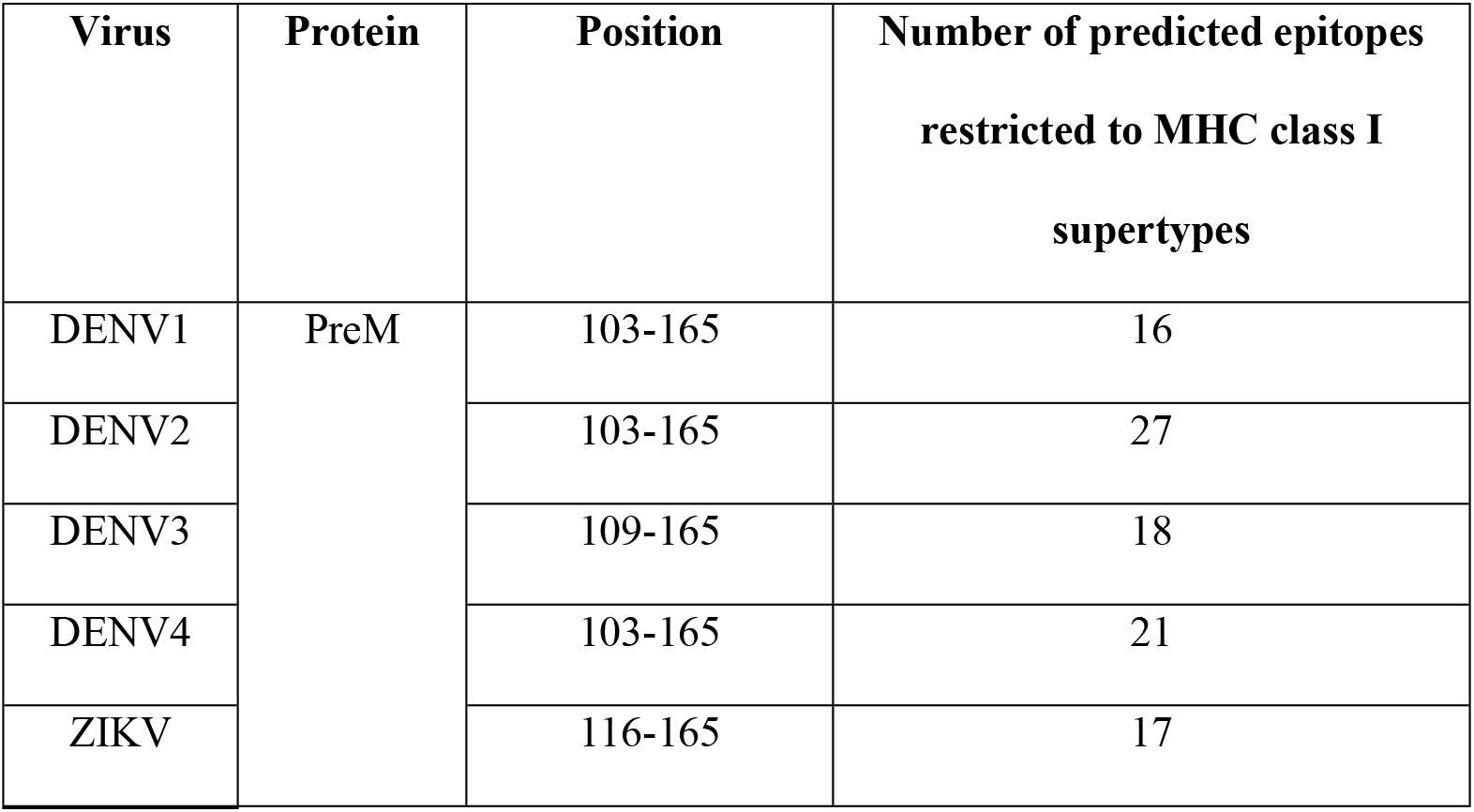
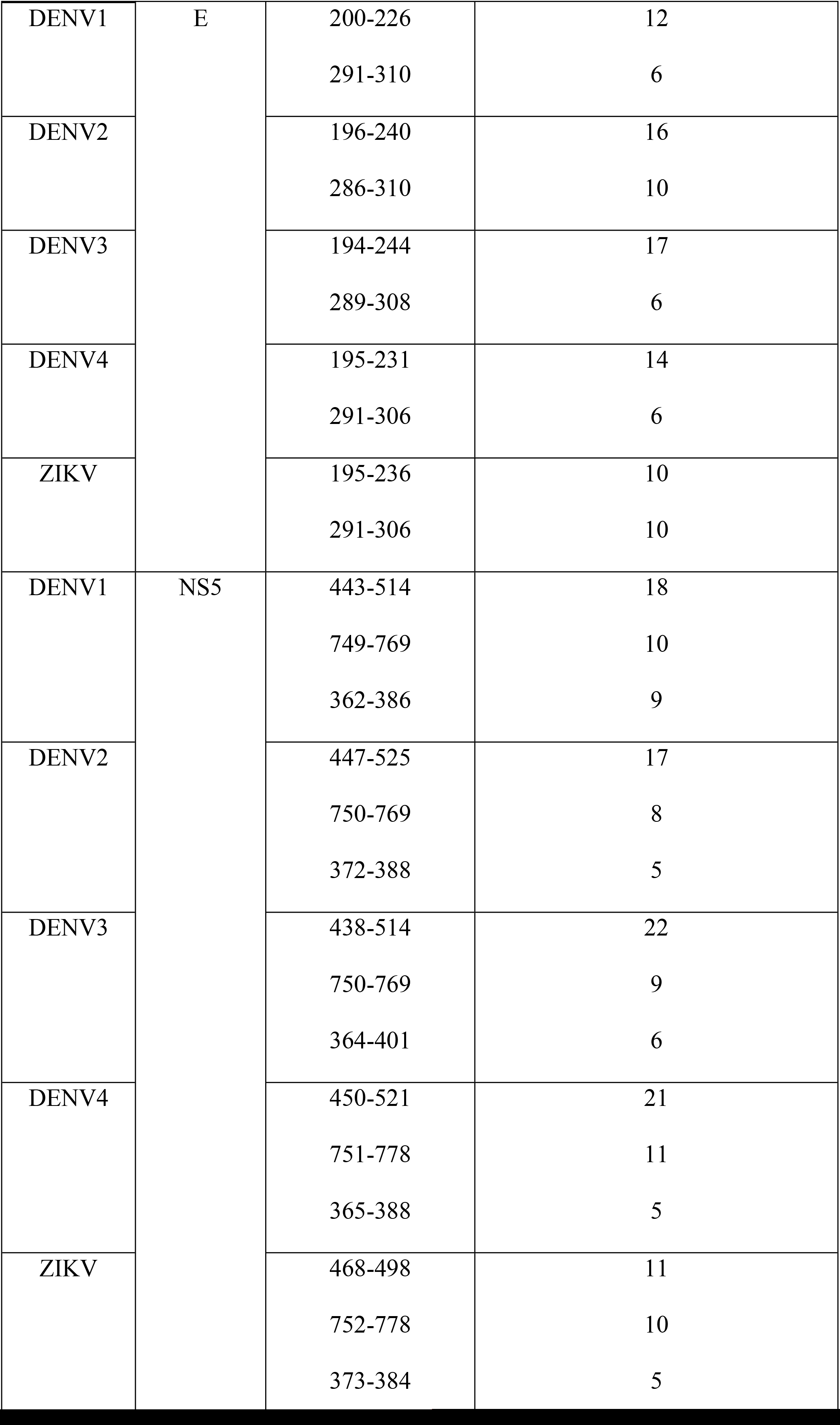
Immunogenic hotspots enriched with multiple MHC class I restricted epitopes from PreM, E and NS5 proteins across all 5 viral species

In contrast, the E protein yielded 2 such dominant immunogenic hotspot stretches **(Table 2)**. Interestingly, E protein, better known for eliciting humoral immune response, gave a 15mer conserved sequence **WLVHKQWFLDLPLPW** within the 200-226 hotspot stretch **(Fig 2(A))**. This conserved sequence has a few amino acid substitutions where L205M, K208R and L212F in DENV3, K210R in DENV2and Q216E, L219H and L221I in ZIKV. Since, the majority of these substitutions are with biochemically similar amino acids and are not in the anchor regions of the epitopes generated from the stretch, this particular hotspot was selected for our MEV candidate. The second hotspot of the E protein did not yield any conserved sequence and therefore was not considered further.

**Fig 2:**
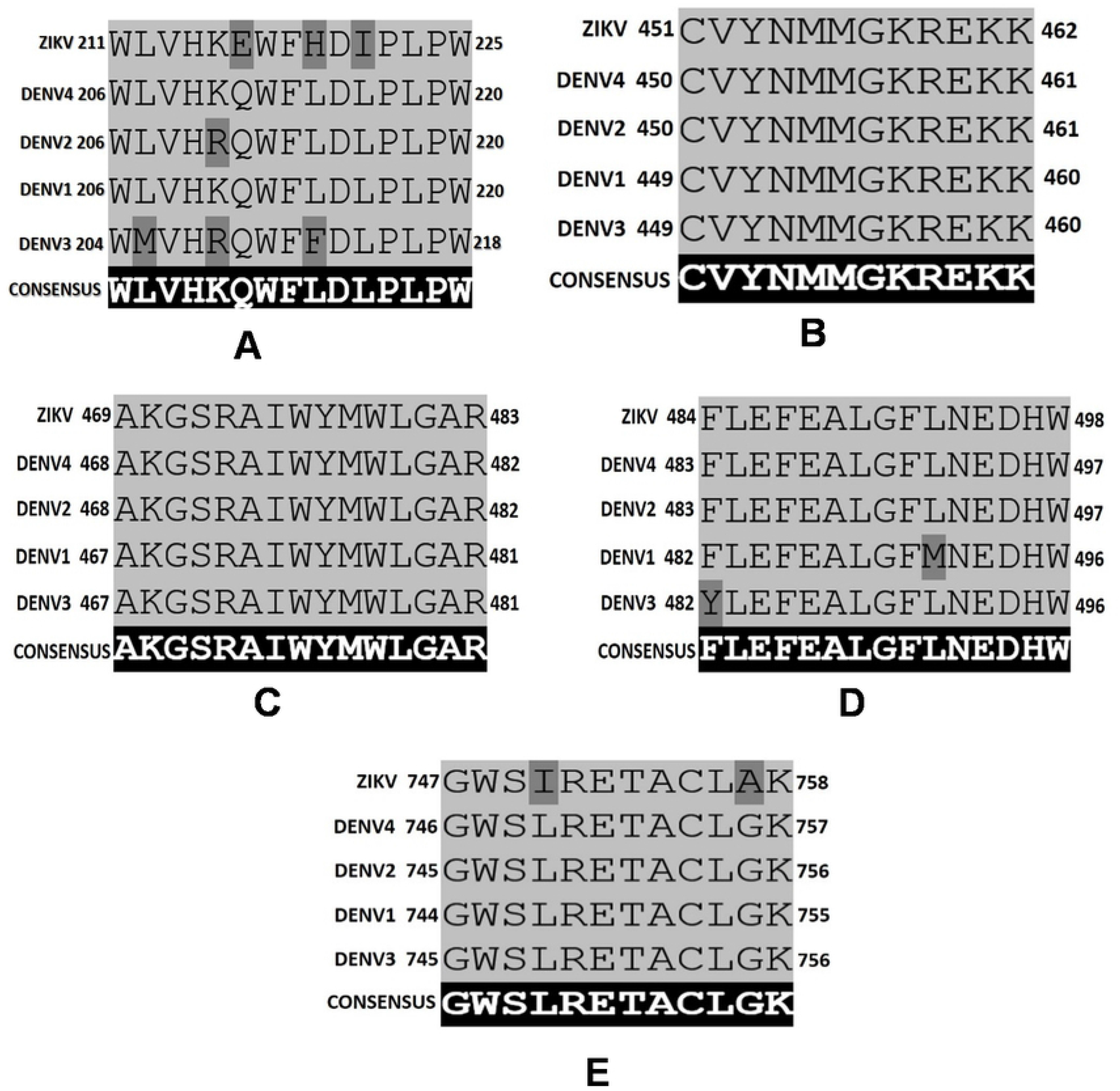
Conserved Immunogenic hotspots identified from the E and NS5 proteins of all DENV serotypes and ZIKV. **(A)** 15mer conserved hotspot from E protein having single position amino acid differences highlighted. **(B)** 12mer sequence from NS5 protein having 100% identity and overlapping epitopes. **(C)** 15mer sequence from NS5 protein having 100% identity and overlapping epitopes. **(D)** Another 15mer hotspot from NS5 protein having single position amino acid differences highlighted. **(E)** 12mer conserved hotspot from NS5 protein having single position amino acid differences highlighted.

NS5, which is known to elicit strong T cell responses, gave three immunogenic hotspot stretches **(Table 2)** that were found in the similar regions across all viral species. Upon investigation of the these stretches, the 450-500 amino acid stretch of NS5 yielded a 12mer **CVYNMMGKREKK** and 15mer **AKGSRAIWYMWLGAR** sequence that was fully conserved containing overlapping epitopes**(Fig 2(B), 2(C))**. Another 15mer sequence **FLEFEALGFLNEDHW** was obtained from the same hotspot, but with two highly-related substitutions (F482Y in DENV3 and L491M in DENV1) **(Fig 2(D))**. A fourth 12mer conserved sequence **GWSLRETACLGK** was mined out from the hotspot stretch 750-769 **(Fig 2(E))**. This sequence had a substitution of two amino acids (L750I and G757A) specifically in ZIKV which are biochemically similar as well. Moreover, these substitutions are not in the anchor region of the epitopes (important for the formation of stable pMHC complexes and antigen presentation by APC) that are generated from these hotspots. Thus, these four “Immunogenic Hotspots” were selected as potential candidates for generation of multi-epitope vaccine.

### Identification of MHC class II restricted epitopes and ‘Immunogenic Hotspot’ from the selected proteins

The MHC class II peptide binding cleft, unlike its class I counterpart, is more open allowing a peptide with higher length and various side chains to interact with the T cell receptor. An optimum of 13 to 17mer peptides is presented by the MHC class II molecules. Therefore, we predicted 15mer epitopes for the MHC class II HLA molecules reference set which has the highest population coverage available on the IEDB server. The predicted epitopes were analyzed to map the immunogenic hotspots. PreM and E proteins each had one hotspot stretch (**Table 3)** that was common for all the viral species. However, upon analysis by multiple sequence alignment, these stretches did not give any conserved peptides for MHC II binding that could be further considered for MEV candidate designing. It is to be noted, that although there was partial overlap of the E protein hotspot with the conserved MHC class I peptide (**WLVHKQWFLDLPLPW**), the overlapping sequence was shorter than the minimum peptide length necessary for MHC class II binding.

**Table 3:**
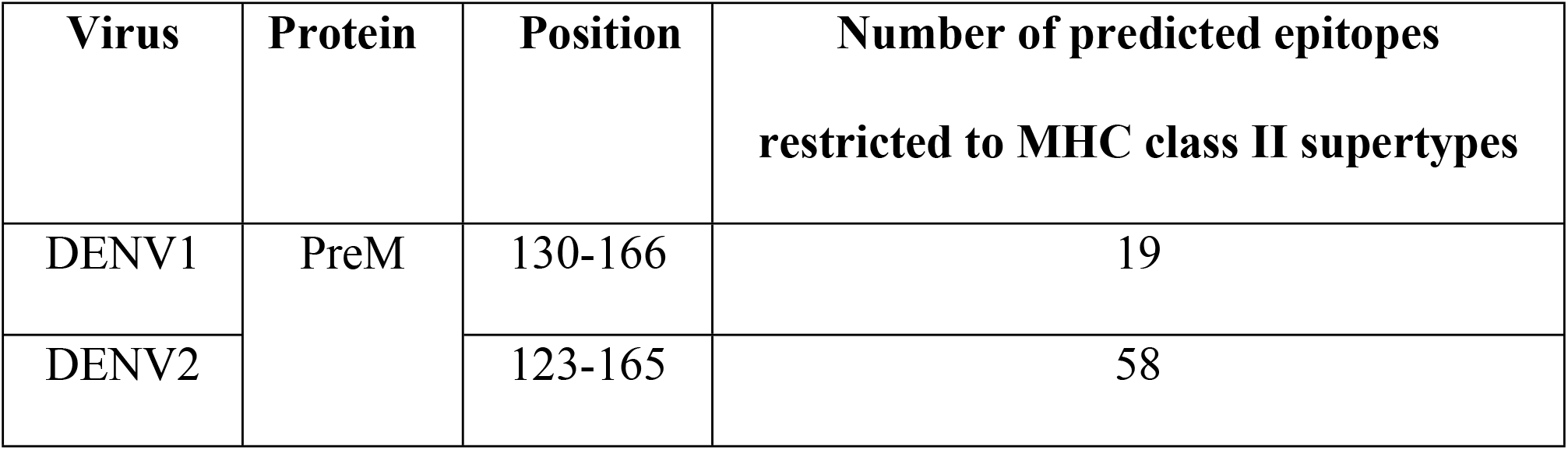
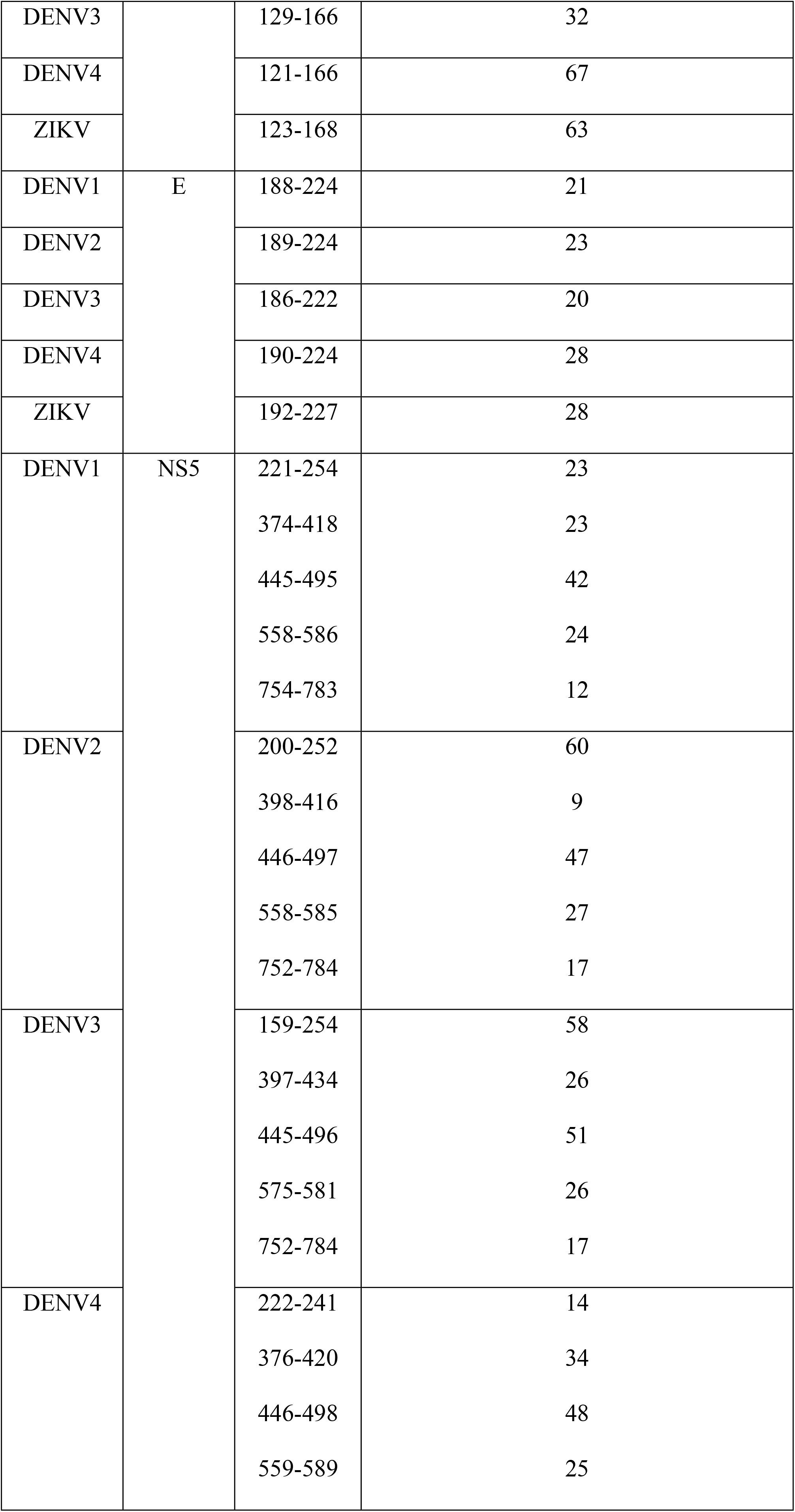
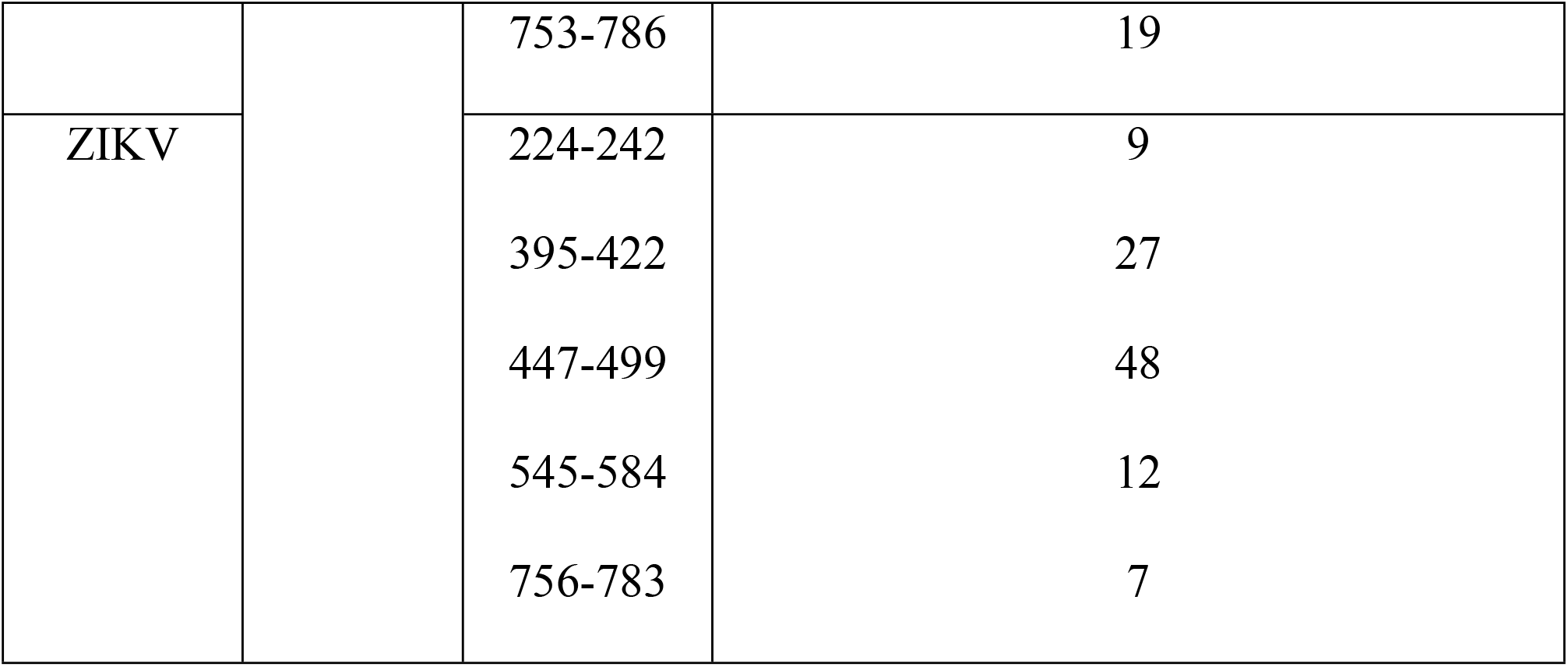
Immunogenic hotspots enriched with multiple MHC class II restricted epitopes from PreM, E and NS5 proteins across all 5 viral species

The NS5 protein gave five immunogenic hotspots, enriched with MHC class II restricted peptides, with three of them coinciding with the hotspot stretches enriched with MHC class I restricted peptides as mentioned in the previous section. Additionally, two hotspot stretches within the 150-250 and 550-590 range having overlapping epitopes was mined out **(Table 3)**. Further investigation revealed a conserved sequence **AKGSRAIWYMWLGARFLEFEALGFLNEDHW** having overlapping epitopes **(Fig 3)**. This 30mer sequence essentially is a combination of the two 15mer hotspots **(Fig 2(C), 2(D))** that were obtained in the case of MHC class I epitopes having identical amino acid replacements. This makes this 30mer hotspot stretch a prime candidate in MEV vaccine design capable of generating a strong CD4^+^ and CD8^+^ T cell response and thereby result in generation of effective memory T cells.

**Fig 3:**
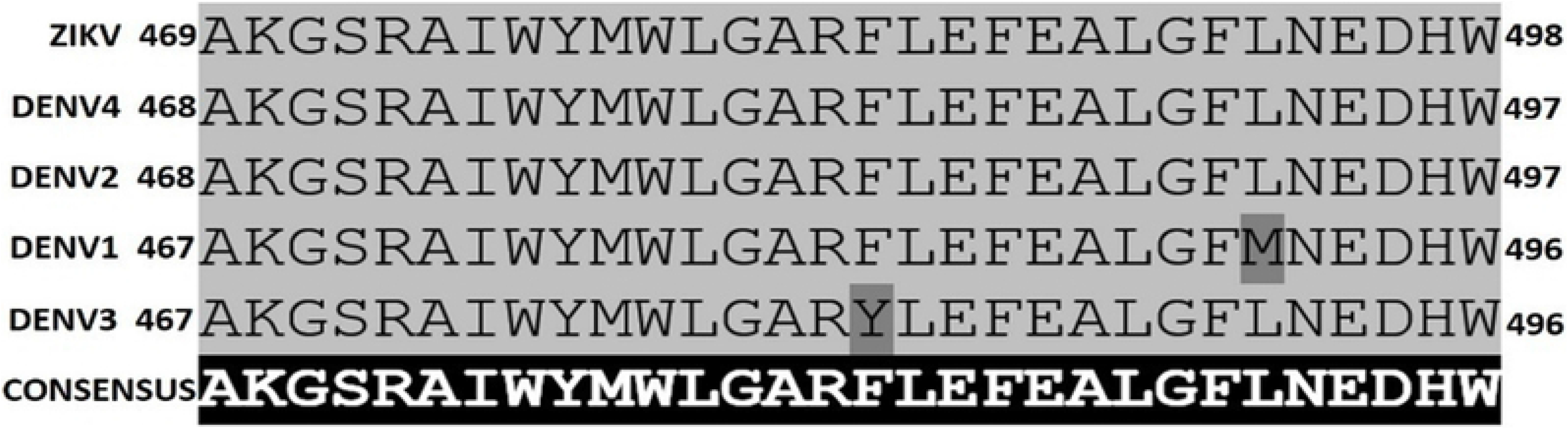
30mer conserved sequence obtained from NS5 protein with single amino acid differences highlighted.

### Prediction of linear B cell epitopes and identification of ‘Immunogenic Hotspot’ from selected proteins

Like T cell epitopes, B cells epitopes were also identified from the candidate proteins. Online epitope prediction tool ABCpred was used for linear B cell epitope prediction of the selected proteins. The 16mer epitopes with threshold value more than 0.75 were selected and further analyzed to identify the immunogenic hotspots.

DENV E and M proteins elicit an initial B cell response that generates neutralizing antibodies against these two proteins. These antibodies against PreM and E proteins have been reported to facilitate secondary infection by a different serotype and increase the viral load via ADE[32,37]. The E protein contains a 15mer immunodominant region at position 97-111 called **fusion peptide loop** (FPL), and antibodies to this highly conserved region frequently show broad flavivirus cross-reactivity resulting in ADE[39,40].Though this region has generated linear epitopes for all DENV serotypes and ZIKV in our studies, but considering its contribution in ADE, it is excluded as an immunogenic hotspot. Furthermore, although an immunodominant region was observed in the E protein, no conserved stretches of considerable length were observed (**S2Table**). Our observation of the conformational B cell epitopes generated from the FPL is described in later sections.

Similar analysis of the PreM protein demonstrated that each serotype of DENV and ZIKV had immunogenic stretches in their sequences that contain multiple linear 16meric B cell epitopes (**S2Table**). However, none of these stretches showed significant overlap. Interestingly, experimental data has reported the generation of at least three antibodies from the PreM protein; one of which is against an identical epitope in ZIKV and DENV serotypes, while the other two are specific to either DENV or ZIKV respectively[41]. The epitope common to both DENV and ZIKV encompasses amino acids derived from position 56-72 of the PreM protein. However, the sequence is not 100% conserved among the viral species, with the conserved sequences being confined to positions 61-70 (**EPdDvDCWCN**) only, with similar amino acid substitutions in position 63 and 65 (denoted with small letter). As discussed earlier about the effect of ADE specifically for these flavivirus infections, the lack of complete conserved sequence of the epitope may lead to greater probability of such incidence;therefore we refrained from utilizing this epitope for the MEV design.

NS5 protein gave maximum number of immunodominant regions for both T cell and linear B cell epitopes (**S2Table**), many of them are in a similar position of the protein sequence. Regions like 440-500, 514-570, and 720-760 have generated B cell epitopes as well as part of both CD4 and CD8 T cell epitopes. The 30mer conserved immunogenic stretch (**AKGSRAIWYMWLGARFLEFEALGFLNEDHW**) (**Fig 3**) previously described also contains multiple linear B cell epitopes. This indicates that this region of NS5 is important for both cell mediated and humoral immune response.

### Conformational B cell epitopes

High resolution (<5Å) and full length three dimensional structures are imperative for structure based computational work including prediction of conformational B cell epitopes. Despite extensive structural analysis of the DENV and ZIKV proteins, there is a lack of fully solved 3-dimensional structures of these proteins in protein data bank (PDB). Most of the available data are either for E protein or the RNA dependent RNA polymerase domain of NS5; both of which have been used for the prediction of conformational B cell epitopes conserved among all serotypes in this study. 3-D structures of either the PreM or M proteins are not available in PDB for all serotypes of the viruses and therefore, these two proteins were excluded from the study. Taking into account the above considerations, conformational B cell epitopes of E and NS5 proteins of DENV serotypes and ZIKV were identified with the help of Discotope 2.0 server using the available 3D structures from PDB.

Only a few regions of the E protein were found to give consistent conformational epitopes. Prominent among them, position 100-102 (GWG) of E protein of ZIKV (PDB ID: 5IRE) and all DENV serotypes except DENV1 (PDB ID: 3G7T) was predicted as epitope. These amino acids are part of the FPL of flavivirus[39,40] which is involved in cross reactivity/ADE among flavivirus members. We further investigated the number of contact residues, which is the number of Cα atoms in the protein within a distance of 10 Å of the residue’s Cα atom, for each epitope. Interestingly, the contact residues of the amino acids constituting the GWG epitope was different for each DENV serotype as well as ZIKV (**S3Table**). Importantly, a similar analysis of the other residues of the FPL, wherever applicable, also demonstrated considerable diversity in the number of contact residues. This could indicate a conformational difference in this region among the viral serotypes of DENV and ZIKV, which may lead to weak interaction of antibodies raised against the FPL of one serotype with the FPL of another,resulting in ADE. Another sequence, 341-346 was predicted as epitopes in all serotypes of DENV, albeit with variable sequence, but not in ZIKV.

Next, we analyzed the conformational epitopes derived from the NS5 protein of ZIKV, DENV2 and DENV3. It is to be noted here that all of the reported structures of the NS5 protein of DENV3 are shorter than actual protein size (899 amino acid residues long), with many of them missing internal sequences. Moreover, due to a complete lack of NS5 structures of DENV1 and DENV4, we could only include the DENV2 and DENV3 NS5 protein structures in our search for B cell conformational epitopes. The conformational epitopes obtained, barring a few, have low sequence homology (**S4Table**). Among them, sequence 14-38 of ZIKV NS5 was found to be overlapping with a common region of predicted conformational epitopes of NS5 protein of DENV 2 and DENV 3 where 23E, 24F, and 27-31 (YKKSG) are conserved. Sequence 107-109 (GPG) was also conserved and predicted as an epitope for both ZIKV and the DENV serotypes. Region 303-318 of ZIKV NS5 also have common embedded conserved epitopes with DENV2 and DENV3, such as 303-305 (TWA), 310Y, 311E. Region 346-369 of ZIKV has a 9mer stretch (VFKEKVDTR) which is also a part of B cell epitope in all DENV serotype under investigation. Similarly, 719-725 and 817-834 has two 4-mer conserved sequences: 721-724 (KDGR) and 828-831(DKTP). However, like the E protein, we further observed the contact residues for each epitope predicted from NS5 (**S4Table**) and likewise found that in majority of the cases, the number of contact residues for each position are different for DENV serotypes and ZIKV. This indicates a difference in conformation of NS5 protein among the serotypes of DENV and ZIKV.

Thus, for both the proteins, conformational B cell epitopes, even when conserved, indicated structural dissimilarities, which could translate into variable affinity of cross-reactive antibodies. Since our aim was to design a single MEV which would be able to induce T and B cell responses without causing ADE, it seemed prudent not to include such conformational epitopes in the candidate.

### Multi-epitope vaccine design

From the above observations and data, it was clear that the MEV candidate must be designed using the conserved immunogenic hotspots derived from E and NS5 protein of DENV and ZIKV that includes MHC class I and MHC class II restricted T cell epitopes and linear B cell epitopes(**Table 4**). To increase the immunogenicity and the probability of uptake by cells, the adjuvant CTxB was included in the N terminus of the MEV. The epitopes were connected with GPGPGPG linker and the adjuvant was connected to the epitopes by EAAAK linker.

**Table 4:**
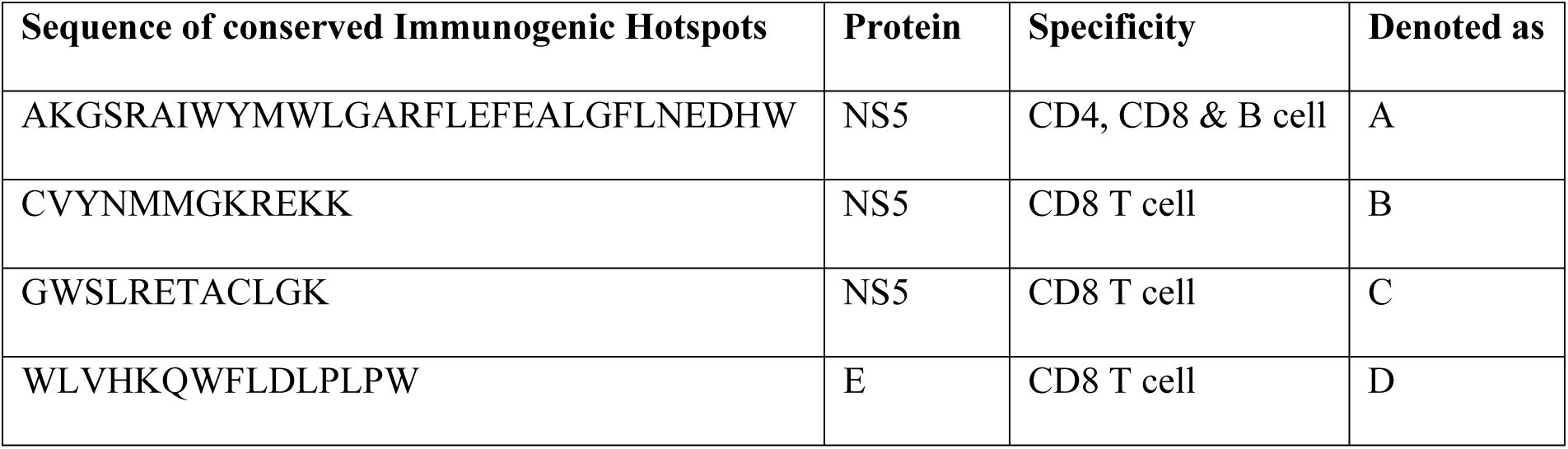
“Immunogenic Hotspots” selected for designing of multi-epitope vaccine from the E and NS5 protein of DENV serotypes and ZIKV

The four immunogenic hotspots were marked alphabetically A, B, C, & D (**Table 4**) and arranged in all possible combinations keeping the CTxB adjuvant at N terminus. The 24 MEV combinations thus generated, were then used for tertiary structure prediction using the trRosetta server. Each combination gave one best predicted structure in PDB format which were named model 1 to 24 and was further evaluated to determine the best candidate(s). The tertiary structures of the terminal adjuvant CTxB is important for the MEV’s internalization into the host APC and thus must remain in its native three-dimensional form. Therefore, the 24 models were aligned with the reported structure of CTxB (PDB ID 5LZG) and the RMSD values were calculated. Models with more than 80% (greater than 82) of the Cα atom aligned with the structure of CTxB and having RMSD less than 1 Å were selected to get the highest possible conserved structures of CTxB. Five models (**8, 9, 13, 19 and 21**) out of 24 that qualified the criteria (**S5 Table**) were further checked for their stability by calculating the Gibbs free energy for folding. This was done using the FoldX server which is used for measurement of protein stability from its 3D structure, by calculating the change in relative free energy for folding of the protein from its unfolded state[42,43]. Except for model 13, the other four models have shown <100 Kcal/mole for folding, indicating a highly stable structure (**S6 Table**). **Fig 4** shows the predicted tertiary structures of the four models (**8, 9, 19 and 21**) as visualized in Pymol. Molecular docking of the individual MEV candidates with TLR4 showed thateach candidate MEV interacts with different regions of the TLR4 and its associated complex MD2 (**Fig 5**). This is indicative of the potential of the selected MEV candidates to activate TLR mediated immune response.These four vaccine candidates were further probed for T and B cell epitope generation using the respective epitope-prediction tools discussed above and it was found that the highly immunogenic T and B cell epitopes have been preserved in these candidates (**S7 Table**).

**Fig 4:**
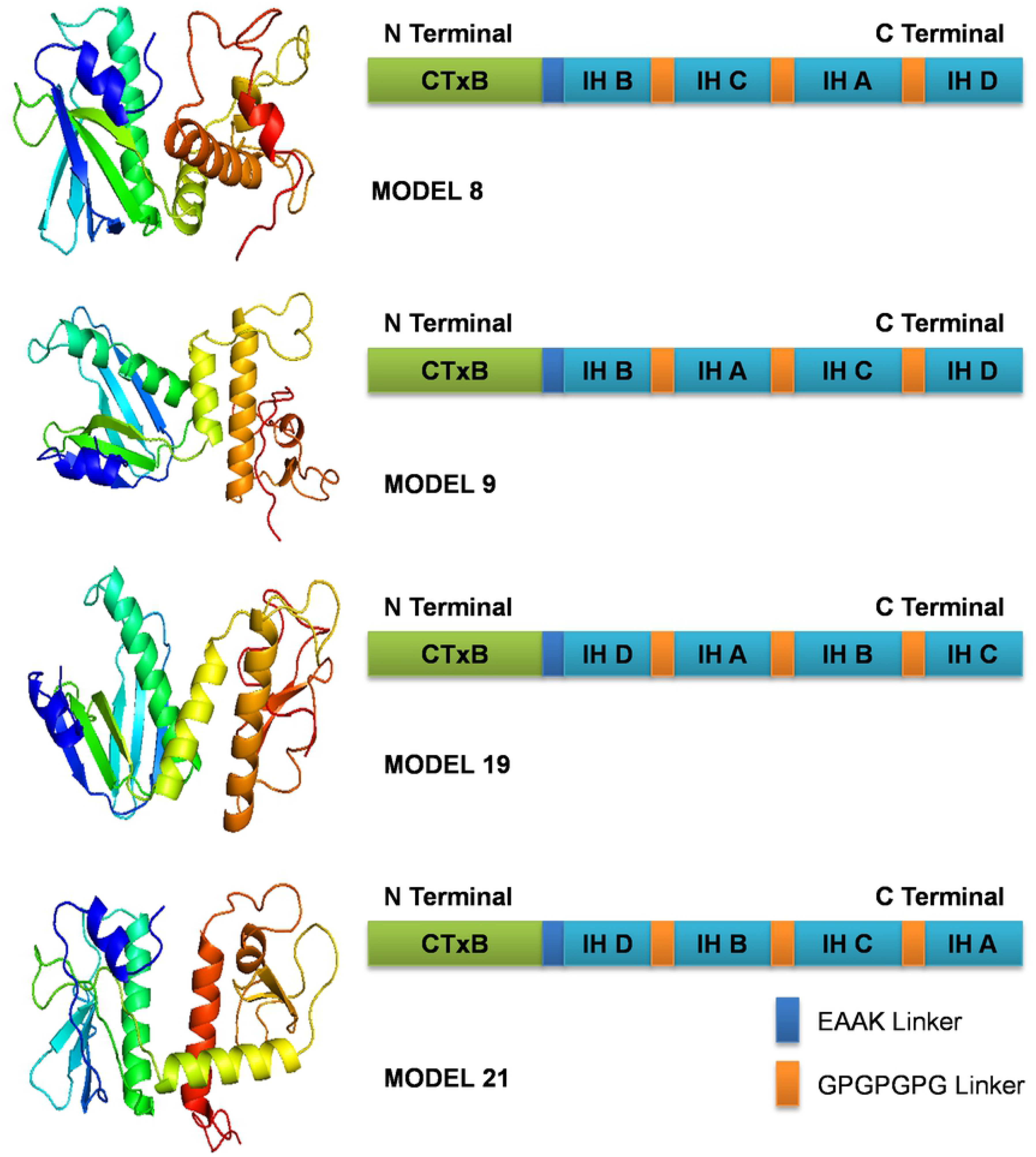
**Predicted tertiary structure of the designed multi-epitope vaccine candidates (8, 9, 19 and 21)** (colored by rainbow from N to C terminus) (IH- Immunogenic Hotspot)

**Fig 5:**
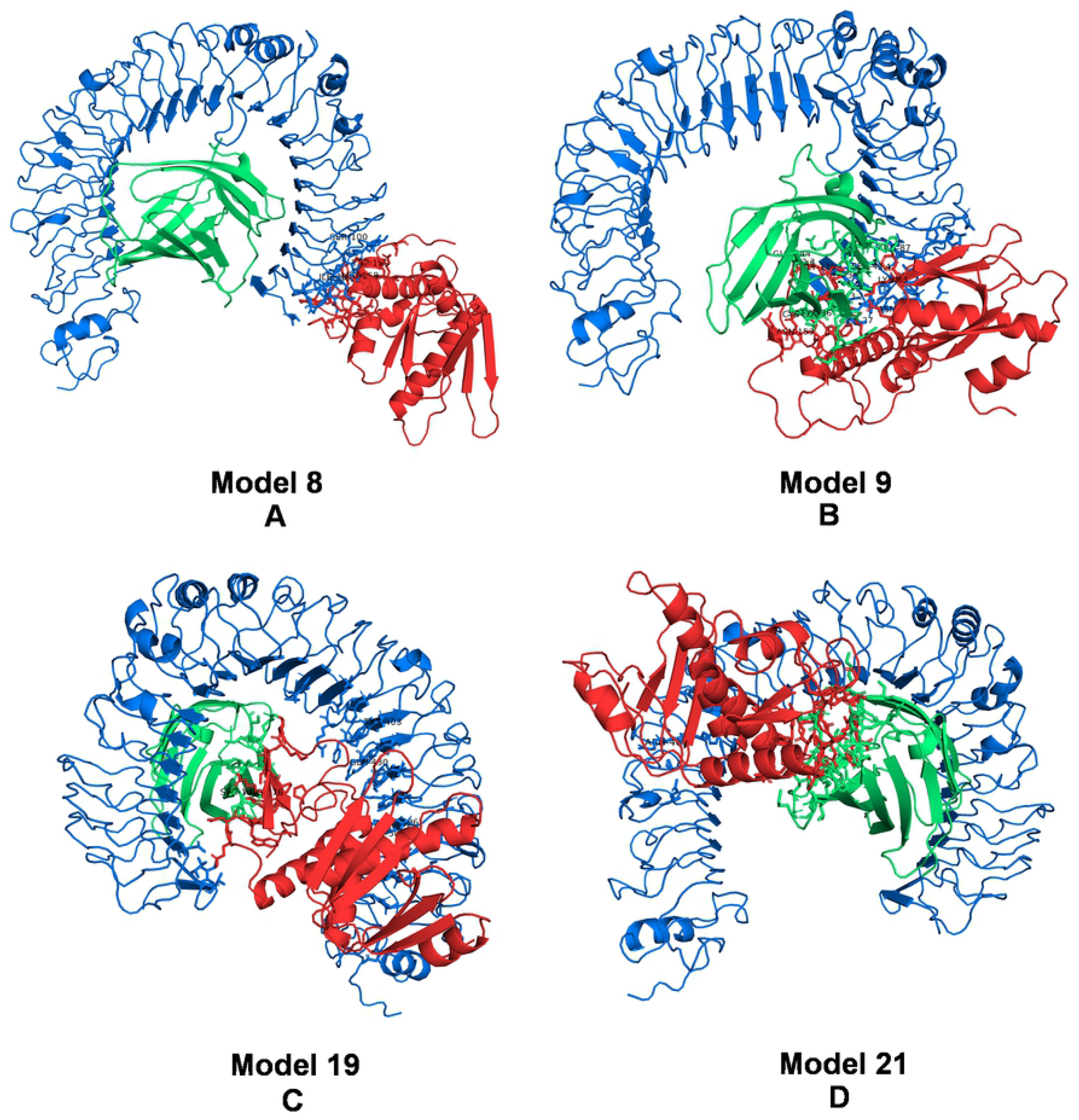
Molecular Docking of MEV candidates with TLR4/MD2 complex. **(A)** Model 8 **(B)** Model 9 **(C)** Model 19 **(D)** Model 21. TLR4 is in blue, MD2 is in green and MEV is in red.

## Conclusion

Conventional MEVs are designed to incorporate multiple MHC class I and MHC class II restricted immunodominant T cell epitopes with the potential to activate multiple T cell clones, which can differentiate into cytotoxic and helper T cells respectively. In addition, such MEVs also include B cell epitopes that can activate B cells leading to their differentiation into antibody-producing plasma cells, which can neutralize the pathogens. One of the most important aspects of MEVs is the exclusion of potentially detrimental parts of antigenic determinants that can otherwise initiate unwanted and harmful immune-responses, like ADE. Especially in the case of flavivirus like DENV and ZIKV, which have a high degree of sequence similarity, ADE is a matter of serious concern, with numerous reports of increased virulence of subsequent heterotypic infections. Therefore, any vaccine made against these viruses, must be specifically checked to exclude any potential ADE, while ensuring the extension of the protection to all serotypes concerned.

Given the close phylogenetic relationship between DENV serotypes and ZIKV, we ventured to design a common vaccine which will not merely incorporate multiple single T and B cell epitopes, but rather include immunogenic stretches or hotspots, rich in multiple HLA supertype-restricted T cell epitopes and B cell epitopes that are common to the pathogens, as well as completely conserved among species. Only four such hotspots could fulfill the stringent criteria, which also specifically excludedthose stretches that contain known ADE-associated epitopes.Notably, in the predicted MEV candidates, only linear B cell epitopes could be included as the predicted conformational B cell epitopes, were not found to be completely conserved sequences among the considered viruses. More importantly, the contact residues of these predicted conformational epitopes varied in number among the serotypes, which could purportedly lead to differences in antibody affinities against these epitopes. Further investigation needs to be done to ascertain the contribution of this observation to ADE.

Furthermore, the designed MEV described here, also incorporates an adjuvant, the CTxB, which can be takenup by cells via the GM1-gangliosides, and has been found to be recognized by TLR4[29], thereby having the ability to initiate TLR mediated signaling pathways that can activate the protective immune response. Lastly, the immunogenic hotspots described in this study include T cell epitopes that are restricted to MHC class I supertypes and MHC class II alleles with maximum population coverage, thus ensuring its efficacy among the global population. In sum, the MEV candidates described here can be considered as a stepping stone towards a common vaccine against DENV and ZIKV.

## Acknowledgment

We would like to thank SERB for funding the project (ECR/2017/002073). We would also like to thank Dr. Dibyendu Samanta and Ms. Kheerthana Duraivelan for their critical inputs which have improved the work significantly.

## Author contributions

**Conceptualization:** Gayatri Mukherjee

**Data Curation:** Dhrubajyoti Mahata, Debangshu Mukherjee, Gayatri Mukherjee

**Formal Analysis:** Dhrubajyoti Mahata, Debangshu Mukherjee, Gayatri Mukherjee

**Funding Acquisition:** Gayatri Mukherjee

**Investigation:** Dhrubajyoti Mahata, Debangshu Mukherjee, Vanshika Malviya, Gayatri Mukherjee

**Methodology:** Dhrubajyoti Mahata, Debangshu Mukherjee, Vanshika Malviya, Gayatri Mukherjee

**Project Administration:** Gayatri Mukherjee

**Resources:** Dhrubajyoti Mahata, Debangshu Mukherjee, Gayatri Mukherjee

**Software:** Dhrubajyoti Mahata, Debangshu Mukherjee

**Supervisor:** Gayatri Mukherjee

**Validation:** Dhrubajyoti Mahata, Debangshu Mukherjee, Gayatri Mukherjee

**Visualization:** Dhrubajyoti Mahata, Debangshu Mukherjee, Gayatri Mukherjee

**Writing- Original Draft Preparation:** Dhrubajyoti Mahata, Debangshu Mukherjee, Gayatri Mukherjee

**Writing- Review and Editing:** Dhrubajyoti Mahata, Debangshu Mukherjee, Gayatri Mukherjee

## Supporting Information

**S1 Table:** Antigenic probabilities of the non-structural proteins as determined by the VaxiJen server.

**S2 Table:** Linear B cell immunogenic hotspots identified from PreM, E and NS5 proteins of all DENV serotypes and ZIKV. The stretches in bold depict common regions to MHC class I and MHC class II restricted immunogenic hotspot.

**S3 Table:** Number of contact residues for thee predicted epitope (101-103) and the rest of the fusion peptide loop region (97-111) of E protein, as predicted by Discotope 2.0 (PDB Id for each protein is put in bracket).

**S4 Table:** Number of contact residues of the conserved epitopes from NS5 protein as predicted by Discotope 2.0(PDB Id for each protein is put in bracket).

**S5 Table:** Alignment result between the known structure of CTxB and the predicted MEV models. The top five candidates are in bold.

**S6 Table:** Gibbs free energy for folding of the selected MEV models.

**S7 Table:** Conserved Epitopes for DENV serotypes and ZIKV as obtained from the MEV models.

## Notes

### Competing Interest Statement

The authors have declared no competing interest.

## References

1. World Health Organization. WHO report on global surveillance of epidemic-prone infectious diseases—dengue and dengue haemorrhagicfever. (2001). [Accessed January 5, 2020] http://www.who.int/csr/resources/publications/dengue/CSR_ISR_2000_1/en/

2. Ioos S, Mallet HP, Leparc Goffart I, Gauthier V, Cardoso T, Herida M. Current Zika virus epidemiology and recent epidemics. Med Mal Infect. 2014; 44(7):302–7. doi: 10.1016/j.medmal.2014.04.008. PMID: 25001879.

3. Rather IA, Lone JB, Bajpai VK, Park YH. Zika Virus Infection during Pregnancy and Congenital Abnormalities. Front Microbiol. 2017; 8:581. doi: 10.3389/fmicb.2017.00581. PMID: 28421065.

4. Rice CM, Lenches EM, Eddy SR, Shin SJ, Sheets RL, Strauss JH. Nucleotide sequence of yellow fever virus: implications for flavivirus gene expression and evolution. Science. 1985; 229(4715):726–33. doi: 10.1126/science.4023707. PMID: 4023707.

5. Wong SS, Poon RW, Wong SC. Zika virus infection-the next wave after dengue? J Formos Med Assoc. 2016; 115(4):226–42. doi: 10.1016/j.jfma.2016.02.002. PMID: 26965962

6. Subramaniam KS, Lant S, Goodwin L, Grifoni A, Weiskopf D, Turtle L. Two Is Better Than One: Evidence for T-Cell Cross-Protection Between Dengue and Zika and Implications on Vaccine Design. Front Immunol. 2020; 11:517. doi: 10.3389/fimmu.2020.00517. PMID: 32269575.

7. Kam YW, Lee CY, Teo TH, Howland SW, Amrun SN, Lum FM, et al. Cross-reactive dengue human monoclonal antibody prevents severe pathologies and death from Zika virus infections. JCI insight. 2017; 2(8):e92428. doi: 10.1172/jci.insight.92428. PMID: 28422757.

8. Stettler K, Beltramello M, Espinosa DA, Graham V, Cassotta A, Bianchi S, et al. Specificity, cross-reactivity, and function of antibodies elicited by Zika virus infection. Science. 2016; 353(6301):823–6. doi: 10.1126/science.aaf8505. PMID: 27417494.

9. Wen J, Shresta S. T Cell Immunity to Zika and Dengue Viral Infections. J Interferon Cytokine Res. 2017; 37(11):475–479. doi: 10.1089/jir.2017.0106. PMID: 29135369.

10. Regla-Nava JA, Elong Ngono A, Viramontes KM, Huynh AT, Wang YT, Nguyen AT, et al. Cross-reactive Dengue virus-specific CD8+ T cells protect against Zika virus during pregnancy. Nat Commun. 2018; 9(1):3042. doi: 10.1038/s41467-018-05458-0. PMID: 30072692.

11. Lim MQ, Kumaran EAP, Tan HC, Lye DC, Leo YS, Ooi EE, et al. Cross-Reactivity and Anti-viral Function of Dengue Capsid and NS3-Specific Memory T Cells Toward Zika Virus. Front Immunol. 2018; 9:2225. doi: 10.3389/fimmu.2018.02225. PMID: 30327651.

12. Dejnirattisai W, Supasa P, Wongwiwat W, Rouvinski A, Barba-Spaeth G, Duangchinda T, et al. Dengue virus sero-cross-reactivity drives antibody-dependent enhancement of infection with zika virus. Nat Immunol. 2016; 17(9):1102–8. doi: 10.1038/ni.3515. Epub 2016 Jun 23. PMID: 27339099.

13. Paul LM, Carlin ER, Jenkins MM, Tan AL, Barcellona CM, Nicholson CO, et al. Dengue virus antibodies enhance Zika virus infection. Clin Transl Immunology. 2016; 5(12):e117. doi: 10.1038/cti.2016.72. PMID: 28090318.

14. Kawiecki AB, Christofferson RC. Zika Virus-Induced Antibody Response Enhances Dengue Virus Serotype 2 Replication In Vitro. J Infect Dis. 2016; 214(9):1357–1360. doi: 10.1093/infdis/jiw377. PMID: 27521359.

15. Halstead SB, O’Rourke EJ. Dengue viruses and mononuclear phagocytes. I. Infection enhancement by non-neutralizing antibody. J Exp Med. 1977; 146(1):201–17. doi: 10.1084/jem.146.1.201. PMID: 406347.

16. Dejnirattisai W, Jumnainsong A, Onsirisakul N, Fitton P, Vasanawathana S, Limpitikul W, et al. Cross-reacting antibodies enhance dengue virus infection in humans. Science. 2010; 328(5979):745–8. doi: 10.1126/science.1185181. PMID: 20448183.

17. Sun J, Du S, Zheng Z, Cheng G, Jin X. Defeat Dengue and Zika Viruses With a One-Two Punch of Vaccine and Vector Blockade. Front Microbiol. 2020; 11:362. doi: 10.3389/fmicb.2020.00362. PMID: 32265852.

18. Halstead SB. Dengvaxia sensitizes seronegatives to vaccine enhanced disease regardless of age. Vaccine. 2017; 35(47):6355–6358. doi: 10.1016/j.vaccine.2017.09.089. PMID: 29029938.

19. Shukla R, Ramasamy V, Shanmugam RK, Ahuja R, Khanna N. Antibody-Dependent Enhancement: A Challenge for Developing a Safe Dengue Vaccine. Front Cell Infect Microbiol. 2020; 10:572681. doi: 10.3389/fcimb.2020.572681. PMID: 33194810.

20. Doytchinova IA, Flower DR. VaxiJen: a server for prediction of protective antigens, tumour antigens and subunit vaccines. BMC Bioinformatics. 2007; 8:4. doi: 10.1186/1471-2105-8-4. PMID: 17207271.

21. Sievers F, Wilm A, Dineen D, Gibson TJ, Karplus K, Li W, et al. Fast, scalable generation of high-quality protein multiple sequence alignments using Clustal Omega. Mol Syst Biol. 2011; 7:539. doi: 10.1038/msb.2011.75. PMID: 21988835.

22. Sidney J, Peters B, Frahm N, Brander C, Sette A. HLA class I supertypes: a revised and updated classification. BMC Immunology. 2008; 9:1. doi: 10.1186/1471-2172-9-1. PMID: 18211710.

23. Greenbaum J, Sidney J, Chung J, Brander C, Peters B, Sette A. Functional classification of class II human leukocyte antigen (HLA) molecules reveals seven different supertypes and a surprising degree of repertoire sharing across supertypes. Immunogenetics. 2011; 63(6):325–35. doi: 10.1007/s00251-011-0513-0. PMID: 21305276.

24. Wang P, Sidney J, Kim Y, Sette A, Lund O, Nielsen M, et al. Peptide binding predictions for HLA DR, DP and DQ molecules. BMC Bioinformatics. 2010; 11:568. doi: 10.1186/1471-2105-11-568. PMID: 21092157.

25. Saha S, Raghava GP. Prediction of continuous B-cell epitopes in an antigen using recurrent neural network. Proteins. 2006; 65(1):40–8. doi: 10.1002/prot.21078. PMID: 16894596.

26. Kringelum JV, Lundegaard C, Lund O, Nielsen M. Reliable B cell epitope predictions: impacts of method development and improved benchmarking.PLoSComput Biol. 2012; 8(12):e1002829. doi: 10.1371/journal.pcbi.1002829. PMID: 23300419.

27. Jia S, Huang X, Li H, Zheng D, Wang L, Qiao X, et al. Immunogenicity evaluation of recombinant Lactobacillus casei W56 expressing bovine viral diarrhea virus E2 protein in conjunction with cholera toxin B subunit as an adjuvant. Microb Cell Fact. 2020; 19(1):186. doi: 10.1186/s12934-020-01449-3. PMID: 33004035.

28. Davod J, Fatemeh DN, Honari H, Hosseini R. Constructing and transient expression of a gene cassette containing edible vaccine elements and shigellosis, anthrax and cholera recombinant antigens in tomato. Mol Biol Rep. 2018; 45(6):2237–2246. doi: 10.1007/s11033-018-4385-3. PMID: 30244396.

29. Phongsisay V, Iizasa E, Hara H, Yoshida H. Evidence for TLR4 and FcRγ-CARD9 activation by cholera toxin B subunit and its direct bindings to TREM2 and LMIR5 receptors. Mol Immunol. 2015; 66(2):463–71. doi: 10.1016/j.molimm.2015.05.008. PMID: 26021803.

30. Yang J, Anishchenko I, Park H, Peng Z, Ovchinnikov S, Baker D. Improved protein structure prediction using predicted interresidue orientations.Proc Natl Acad Sci U S A. 2020; 117(3):1496–1503. doi: 10.1073/pnas.1914677117. PMID: 31896580.

31. Pierce BG, Wiehe K, Hwang H, Kim BH, Vreven T, Weng Z. ZDOCK server: interactive docking prediction of protein-protein complexes and symmetric multimers. Bioinformatics. 2014;30(12):1771–3. doi: 10.1093/bioinformatics/btu097. PMID: 24532726.

32. Valdés K, Alvarez M, Pupo M, Vázquez S, Rodríguez R, Guzmán MG. Human Dengue antibodies against structural and nonstructural proteins. Clin Diagn Lab Immunol. 2000; 7(5):856–7. doi: 10.1128/CDLI.7.5.856-857.2000. PMID: 10973471.

33. Lai CY, Tsai WY, Lin SR, Kao CL, Hu HP, King CC, et al. Antibodies to envelope glycoprotein of dengue virus during the natural course of infection are predominantly cross-reactive and recognize epitopes containing highly conserved residues at the fusion loop of domain II. J Virol. 2008; 82(13):6631–43. doi: 10.1128/JVI.00316-08. PMID: 18448542.

34. Dai L, Song J, Lu X, Deng YQ, Musyoki AM, Cheng H, et al. Structures of the Zika Virus Envelope Protein and Its Complex with a Flavivirus Broadly Protective AntibodyCell Host Microbe. 2016; 19(5):696–704. doi: 10.1016/j.chom.2016.04.013. PMID: 27158114.

35. Rivino L, Kumaran EA, Jovanovic V, Nadua K, Teo EW, Pang SW, et al. Differential targeting of viral components by CD4+ versus CD8+ T lymphocytes in dengue virus infection. J Virol. 2013; 87(5):2693–706. doi: 10.1128/JVI.02675-12. PMID: 23255803.

36. Junjhon J, Edwards TJ, Utaipat U, Bowman VD, Holdaway HA, Zhang W, et al. Influence of pr-M cleavage on the heterogeneity of extracellular dengue virus particles. J Virol. 2010; 84(16):8353–8. doi: 10.1128/JVI.00696-10. PMID: 20519400.

37. Rodenhuis-Zybert IA, van der Schaar HM, da Silva Voorham JM, van der Ende-Metselaar H, Lei HY, Wilschut J, et al. Immature dengue virus: a veiled pathogen? PLoSPathog. 2010; 6(1):e1000718. doi: 10.1371/journal.ppat.1000718. PMID: 20062797.

38. Frank SA. Immunology and Evolution of Infectious Disease. Princeton (NJ): Princeton University Press; 2002. PMID: 20821852.

39. Allison SL, Schalich J, Stiasny K, Mandl CW, Heinz FX. Mutational evidence for an internal fusion peptide in flavivirus envelope protein E. J Virol. 2001; 75(9):4268–75. doi: 10.1128/JVI.75.9.4268-4275.2001. PMID: 11287576.

40. Stiasny K, Kiermayr S, Holzmann H, Heinz FX. Cryptic properties of a cluster of dominant flavivirus cross-reactive antigenic sites. J Virol. 2006; 80(19):9557–68. doi: 10.1128/JVI.00080-06. PMID: 16973559.

41. Amrun SN, Yee WX, Abu Bakar F, Lee B, Kam YW, Lum FM, Tan JJ, Lim VW, Watthanaworawit W, Ling C, Nosten F, Renia L, Leo YS, Ng LF. Erratum: Novel differential linear B-cell epitopes to identify Zika and dengue virus infections in patients. Clin Transl Immunology. 2020; 9(2):e01118. doi: 10.1002/cti2.1118. PMID: 32099653.

42. Schymkowitz J, Borg J, Stricher F, Nys R, Rousseau F, Serrano L. The FoldX web server: an online force field. Nucleic Acids Res. 2005; 33(Web Server issue):W382–8. doi: 10.1093/nar/gki387. PMID: 15980494.

43. Buß O, Rudat J, Ochsenreither K. FoldX as Protein Engineering Tool: Better Than Random Based Approaches? Comput Struct Biotechnol J. 2018; 16:25–33. doi: 10.1016/j.csbj.2018.01.002. PMID: 30275935.

